# Acoustic frequency-dependent physical mechanism of sub-MHz ultrasound neurostimulation

**DOI:** 10.1101/2021.09.11.458049

**Authors:** Haixiao Fan, Kenta Shimba, Ayumu Ishijima, Kenya Sasaoka, Tsuyoshi Takahashi, Chih-Hsiang Chang, Yasuhiko Jimbo, Takashi Azuma, Shu Takagi

## Abstract

Ultrasound is an innovative physical modality allowing non-invasive and reversible modulation of neural circuit activity in the brain with high spatial resolution. Despite growing interest in clinical applications, the safe and effective use of ultrasound neuromodulation has been limited by a lack of understanding of the physical mechanisms underlying its effects. Here, we demonstrate acoustic frequency-dependent physical effects that underlie ultrasound neuromodulation, where cavitation and radiation forces are the dominant sources of low- and high-frequency stimulation, respectively. We used 39.5 kHz and 500 kHz acoustic frequencies to stimulate cultured neural and glial cells, excised from rat cortex, to study acoustic frequency-dependent neural responses. We demonstrate increased evoked responses due to increased cavitation activity at the 39.5 kHz acoustic frequency. In contrast, notable cavitation activity was not detected at 500 kHz despite detection of evoked responses. Our work highlights the dependence of ultrasound neuromodulation on acoustic frequencies, with different physical effects underlying cell responses to low and high sub-MHz acoustic frequency ranges.

## 1. Introduction

Non-invasive neuromodulation has increased our understanding of complex neural functions and has advanced neuromodulation technologies toward clinical applications [1]. However, conventional non-invasive methods such as electrical [2] and magnetic techniques [3] are limited by low spatial targeting capability. Although optogenetics enables single-cell neurostimulation [4], [5], modulation targets are limited to shallow regions of biological tissues due to the scattering of visible light. Furthermore, genetic modifications are often required for optogenetic techniques, which is not acceptable in humans. Despite advancements in modulating neural circuit activity and brain function, most of these methods are applicable only in animal models. The use of ultrasound in previous decades led to a new era in neuromodulation, demonstrating its non-invasiveness and precise stimulation of targeted deep tissue sites [6], [7]. Technological developments of ultrasound neuromodulation have expanded in recent years, with efficacy reported in various animal models [8-16] and human subjects using sub-MHz ultrasound via transcranial approaches [17], [18].

Despite growing interest in neurostimulation and its therapeutic applications, the underlying physical mechanisms remain unclear. Understanding of the underlying mechanism is critical for ensuring safe and effective ultrasound neuromodulation. The two major posited mechanisms for transferring ultrasound energy to tissues are thermal and mechanical effects [19]. A number of studies have reported that ultrasound can modulate neuronal activity without significant heating of tissues [20-22]. The recent leading candidates for the physical mechanisms underlying neuromodulation are mechanical effects originating from cavitation or radiation forces to alter bilayer permeability [23], membrane capacitance [24], mechanosensitive ion channels [6], [25] and morphological changes including neuronal retraction and cell body shrinkage [26]. It should be considered that acoustic frequency dependence is inherent to both cavitation and radiation forces. However, comprehensive insights are still lacking to explain the contribution of acoustic frequency dependence on neuromodulation efficacy. Several *in vivo* studies using sub-MHz ultrasound have reported increased neural responses with decreasing acoustic frequency. Contrarily, *in vitro* studies using ∼10 MHz ultrasound have demonstrated increased neural responses with increasing acoustic frequency. Further studies of neuromodulation are required to clarify these contradictory findings.

Here, we demonstrate acoustic frequency-dependent physical effects that underlie ultrasound neuromodulation, where cavitation and radiation forces are the dominant sources of low- and high-frequency stimulation, respectively **(Fig. 1)**. In the process of stable or unstable cavitation, bubble-oscillations or micro-streaming can deform cellular membranes [27]. The radiation force has also been shown to induce mechanical displacement of cells and tissues [28], [29]. Both physical effects are dependent on acoustic frequencies, with occurrence probability of cavitation decreasing with increasing frequencies [30], and the radiation force increasing with increasing frequencies [31]. Accordingly, we used 39.5 kHz and 500 kHz acoustic frequencies to stimulate cells cultured from rat cortex to study frequency-dependent cell responses. Intracellular Ca^2+^ levels were measured with acoustic spectra to detect cavitation. We found that cavitation impacts neuron activity at 39.5 kHz acoustic frequency, whereas cavitation activity was not confirmed at 500 kHz despite detection of evoked neural responses. These results increase the understanding of acoustic frequency-dependent mechanisms underlying ultrasound neuromodulation, where different physical effects activate neural responses in low and high acoustic frequency ranges.

**Fig. 1.**
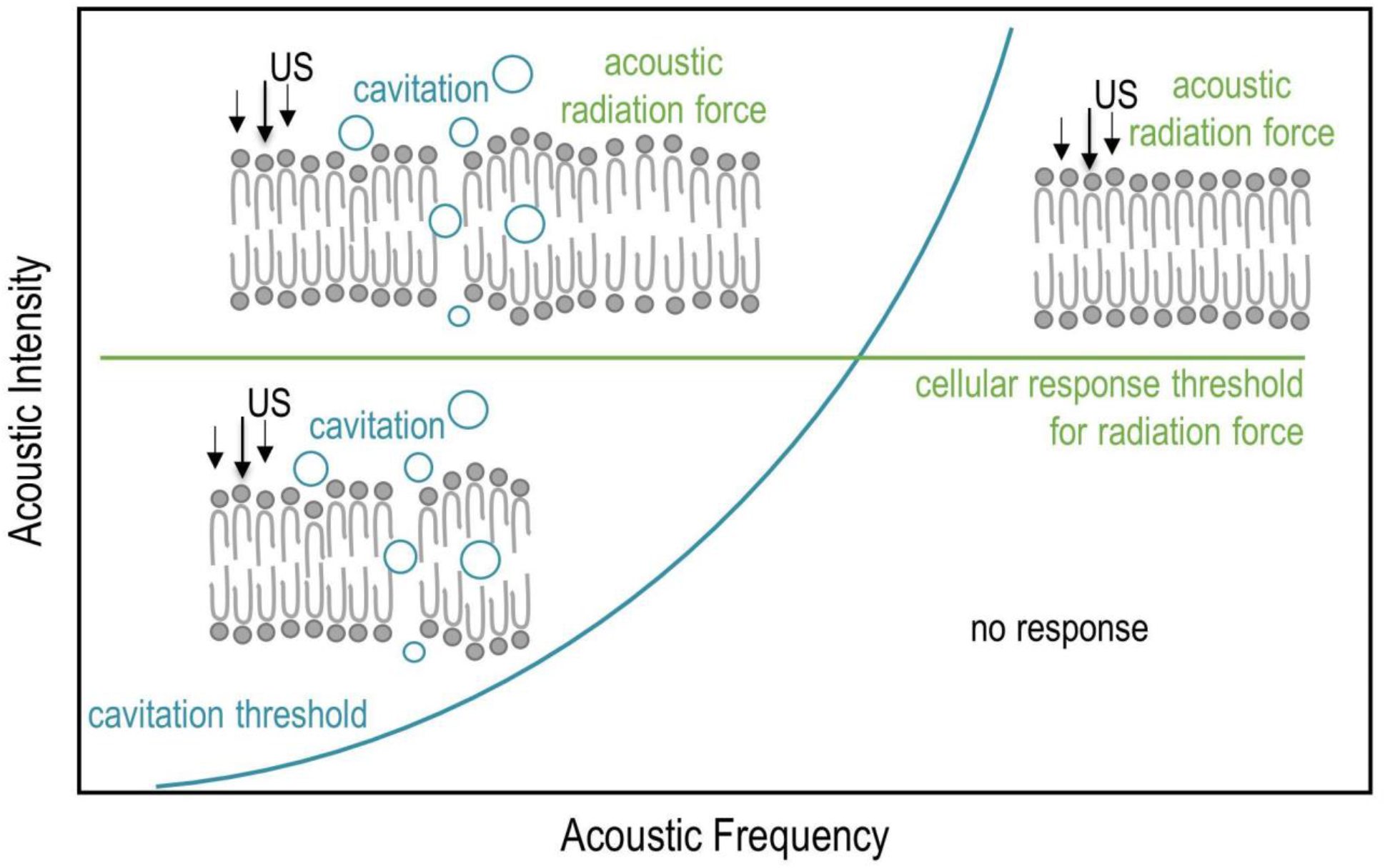
Proposed model of acoustic frequency-dependent physical effect of ultrasound neuromodulation. Cavitation is less likely to occur at higher acoustic frequencies whereas radiation forces increase with increasing acoustic frequencies. The cavitation threshold is below the cellular response threshold for radiation forces at low-frequency ranges, hence cavitation can induce cellular responses with low acoustic intensity. At high-frequency ranges, the cavitation threshold is below the cellular response threshold for the radiation force. Therefore, the radiation force can induce cellular responses at acoustic intensities below the cavitation threshold.

## 2. Materials and Methods

### 2.1 Cell culture

Studying the mechanisms underlying ultrasound neuromodulation *in vivo* is challenging due to off-target auditory ultrasound responses [32], [33]. We therefore used primary cell cultures in the present study. All animals were maintained following the regulations set by The University of Tokyo, and all animal experiments were conducted following the institutional guidelines with the approved protocol. Cortical tissues were dissected from embryonic day 19 Wistar rat embryos (Oriental Yeast, Tokyo, JP). Cells were maintained in neurobasal medium (21103049, Invitrogen, US) supplemented with 1× B27 (17504044, Thermo Fisher, US), 2 mM GlutaMax (35050061, Invitrogen, US) and 1% penicillin-streptomycin (15140122, Thermo Fisher, US). Before seeding, culture dishes were coated with polyethylenimine (P3143, Sigma, US)/laminin (23017015, Thermo Fisher, US). After removing the coating solution, cells were seeded at an initial cell density of 1000 cells/mm^2^. Cells were cultured inside a neurobasal medium-based conditioned medium from mature cortical cells to support survival and growth of the newly-seeded cortical culture. After the first week of culture, half of the medium was exchanged every three days. All ultrasound stimulation experiments were performed at 28 days after seeding.

### 2.2 Acoustic pressure profile measurement

The acoustic pressure profile of the ultrasound transducers was measured inside a degassed water tank using a needle hydrophone (Type 80-0.5-4.0, Imotec Messtechnik, DE). The hydrophone position was stepped with a 1-mm pitch by a three-axis motorised stage (ALS-305 -CM, ALZ-305- CM, ALS-604-E1P, Chuo Precision Industrial, JP) and a stage controller (QT- CM2, Chuo Precision Industrial, JP). Voltage traces were recorded by a digital oscilloscope (HDO4024, Teledyne LeCroy, US).

### 2.3 Ultrasound stimulation

An arbitrary waveform generator (WF1974, NF Corporation, JP) generated the desired number of cycles of a sinusoidal waveform. The generator output was amplified by an amplifier (2100L, Electronics & Innovation, US) and used to drive the transducer. Two types of unfocused ultrasound transducers, 39.5 kHz (15 mm aperture, FBL40152SSF-FC, Fuji Ceramics Corporation, JP) and 500 kHz (13.5 mm aperture, 0.5K1OI, Japan Probe, JP), were used to generate ultrasound. The transducers were driven by the amplified 39.5 kHz and 500 kHz sinusoidal waves with a pulse duration of 41.7 ms. Acoustic intensity-dependent responses were measured by changing the delivered spatial-peak pulse average intensity (I_SPPA_) to 0.03– 0.49 W/cm^2^ for 39.5 kHz and 0.19–1.11 W/cm^2^ for 500 kHz. The time interval for changing the acoustic power was set as 3 min to avoid heating of transducer. The distance between the ultrasound transducer and the sample was set as 3 mm and 5 mm for 39.5 kHz and 500 kHz, respectively, to avoid near-field interference [20].

### 2.4 Acoustic signal recording

Acoustic signals during the ultrasound stimulation were recorded by a needle hydrophone (Type 80-0.5-4.0, Imotec Messtechnik, DE) connected to a digital oscilloscope (HDO4024, Teledyne LeCroy, US). The distances between the hydrophone, ultrasound transducer and sample were fixed with a custom-made jig. The second harmonic frequency and broadband noise of the recorded signals were analysed to detect stable and inertial cavitation during ultrasound stimulation, respectively [34-37]. Spectral amplitude *I* was normalised by the amplitude at fundamental frequency *I*_*0*_. The normalised amplitude *I* / *I*_*0*_ at 1.8*f*_*0*_–2.2*f*_*0*_ and 1.2*f*_*0*_–1.8*f*_*0*_ was used to calculate the spectral energy of second harmonic and broadband noise, respectively (*f*_*0*_ denotes fundamental frequency).

### 2.5 Ca^2+^ fluorescence imaging

Changes in intracellular Ca^2+^ concentrations were measured using a fluorescence microscope (IX71, Olympus, JP) equipped with a laser source (Sapphire LP, Coherent, US), a confocal scanner unit (CSU-X1, Yokogawa, JP), an image sensor (C8800, Hamamatsu Photonics, JP) and a stage incubator (INU-TIZ-F1, Tokai Hit, JP). Immediately before ultrasound stimulation experiments, cells were incubated with 3 μL of 10 mM Fluo-8 AM solution (21082, AAT Bioquest, US), 1.2 μL of 10% (w/v), non-ionic detergent Pluronic F-127 solution (P6866, Thermo Fisher, US) and 600 μL recording buffer containing 140 mM NaCl, 5 mM KCl, 1 mM MgCl_2_, 5.5 mM glucose, 2 mM CaCl_2_ and 10 mM HEPES for 30 min to visualise changes in intracellular Ca^2+^ concentrations. After washing cells with recording buffer twice, cells were incubated for 15 min and imaged with a fluorescence microscope. Fluorescence intensities before the onset of ultrasound stimulation were used as a baseline for analysing responses upon ultrasound stimulation. Calcium signals in a time window between 0 s (onset of ultrasound) and 1 s were used to calculate response delay. An increase in fluorescence intensity before and after ultrasound stimulation was normalised using the resting value. Fluorescence curves were fitted with the four parameters logistic regression method (R^2^ > 0.95). We defined response delay as the time when the amplitude of the fitted fluorescence curve increased to 0.2% [38]. Ultrasound-activated cells were defined as the amplitude of fluorescence increased to 0.2% compared with the baseline. Response rate per dish was defined as the proportion of activated cells out of all recorded cells.

### 2.6 Radiation force calculation

The radiation force *F* (N/m^3^) generated in ultrasound field was estimated by using following equation [39]:

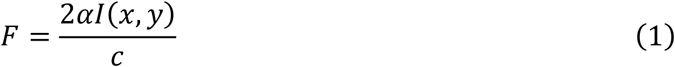

where *α* (Np/m) is the absorption coefficient, *I* (W/m^2^) is the acoustic intensity, and *c* (m/s) is the sound speed. The absorption coefficient *α* can be expressed as:

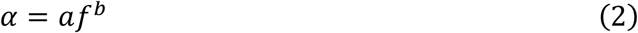

where *f* (MHz) is the acoustic frequency, *a* and *b* are constants varying in different tissues. Here, we assumed the sound speed to be 1560 m/s [40], *a* to be 0.024 Np/cm/MHz^-b^, and *b* to be 1.18 for brain tissue [41].

### 2.7 Statistical analysis

The two-sample two-tailed *t*-test was used to evaluate significance between different experimental groups.

## 3. Results and Discussion

### 3.1 Intracellular Ca^2+^ elevation evoked by sub-MHz ultrasound stimulation

Frequency-dependent neural responses of cells cultured from rat cortex to ultrasound stimulation were studied by recording neural calcium responses with acoustic spectra to detect cavitation **(Fig. 2a, b)**. Transducers delivered uniform acoustic pressure throughout the field of view of recorded fluorescence images **(Fig. 2c)**.

**Fig. 2.**
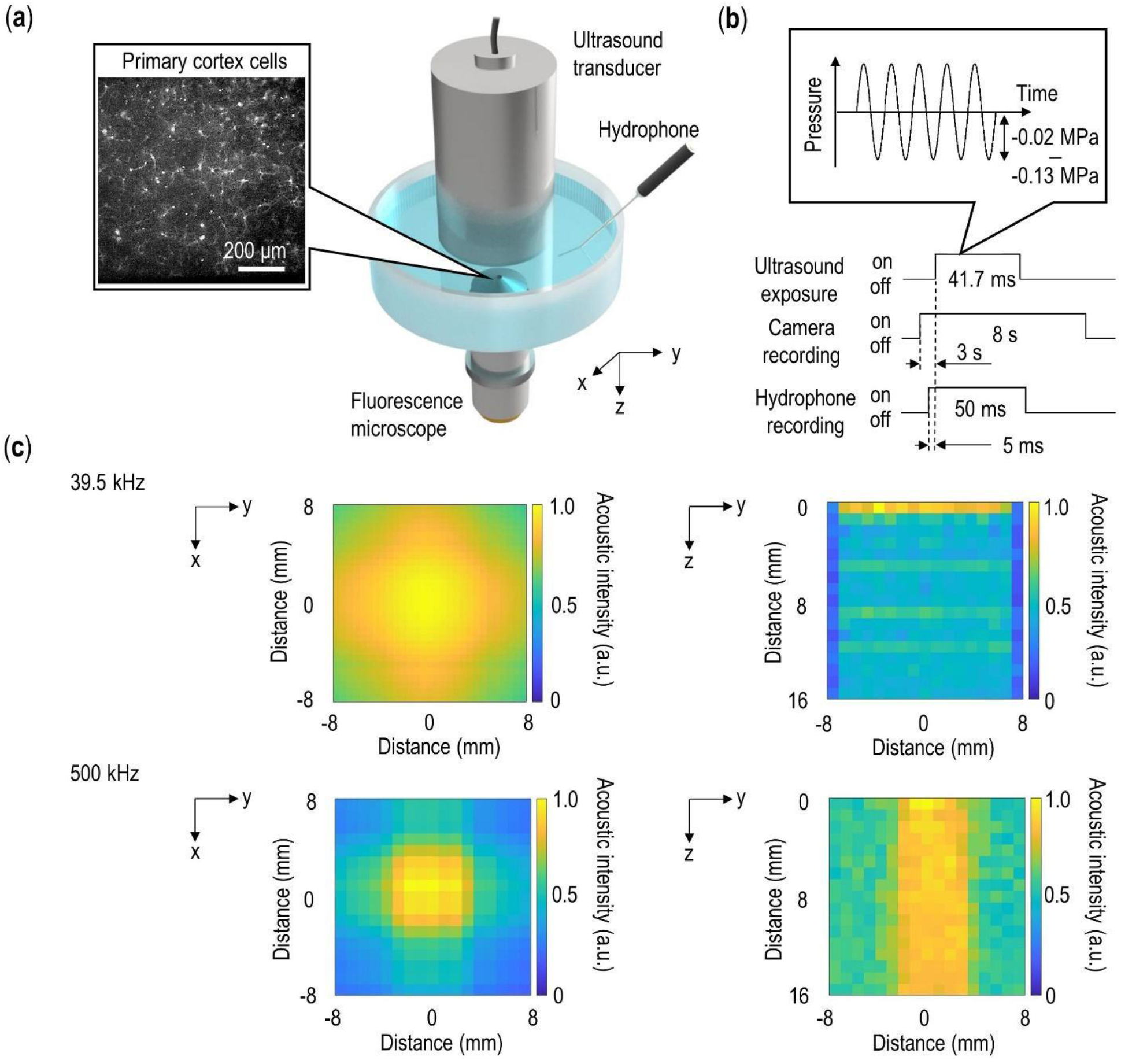
Ultrasound neurostimulation setup. **(a)** The ultrasound transducer was immersed inside the medium in order to expose cultured neurons to ultrasound. Acoustic signals were recorded with the hydrophone to detect the occurrence of cavitation. Changes in cellular Ca^2+^ concentration were measured by recorded cellular responses to ultrasound exposure using a fluorescence microscope. **(b)** Timing schematic for the experimental setup. **(c)** The acoustic intensity distribution of 39.5 kHz (top) and 500 kHz (bottom) in the lateral direction at a distance of 3 mm and 5 mm from the transducer surface and the axial direction, respectively.

Ultrasound stimulation with 39.5 kHz and 500 kHz evoked Ca^2+^ concentration changes inside neurons **(Fig. 3a, Supplementary Video 1 and 2)**. We investigated fluorescence changes under different acoustic intensities to determine the association between acoustic intensity and cell responses to ultrasound stimulation **(Fig. 3b, c)**. Evoked responses were measured by analysing transient changes in Ca^2+^ concentration in response to ultrasound stimulation. Cortical cells were activated by 39.5 kHz ultrasound stimulation with lower acoustic intensities compared to 500 kHz ultrasound stimulation. This trend was also presented in the relationship between response rate and acoustic intensity **(Fig. 4b, c)** and it was similar to previously reported in vivo neuromodulation studies using sub-MHz ultrasound, where higher intensities were required for high-frequency stimulation [39]. Moreover, activated cells showed approximately 150 ms delay in calcium responses, which was reduced by increasing ultrasound intensity **(Fig. 3d, e)**. This delayed onset is typical of neural responses to low intensity ultrasound [38], where mechanical effects slowly deplete ion gradients with subsequent membrane depolarisation [1].

**Fig. 3.**
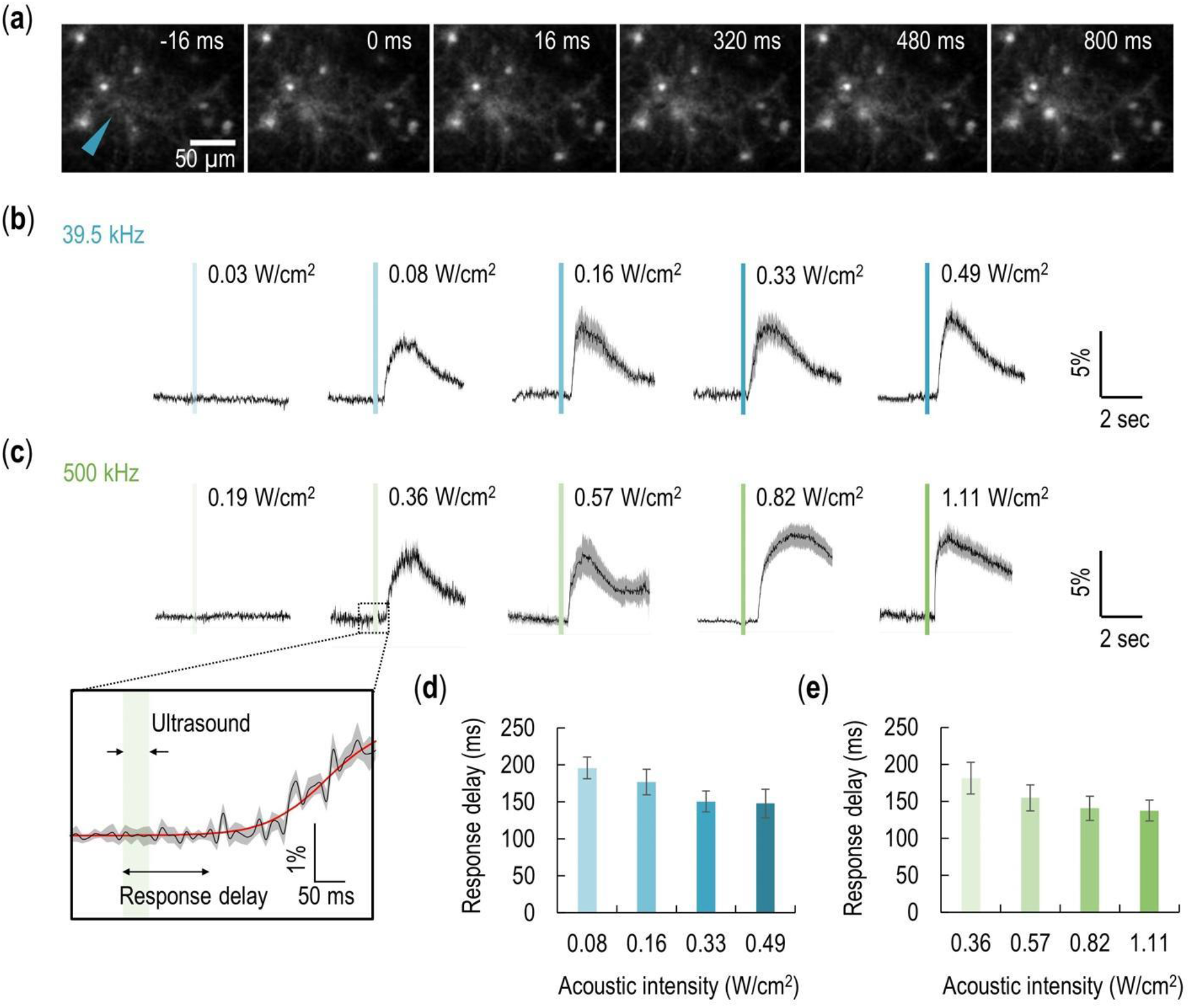
Calcium responses of ultrasound-stimulated cells. **(a)** Representative time-lapse fluorescence images of cultured cortical cells recorded at a frame rate of 60 fps during and after ultrasound stimulation (39.5 kHz, 0.33 W/cm^2^, 41.7 ms pulse duration). **(b)** Ca^2+^ fluorescent transient curves of ultrasound-stimulated cells at acoustic frequencies of 39.5 kHz and **(c)** 500 kHz (N = 6 dishes). Inset shows a representative cell response delay to the onset of ultrasound stimulation with a fitted curve shown as the red line. **(d)** Response delay time for 39.5 kHz and **(e)** 500 kHz ultrasound stimulation. Shaded areas and error bars represent standard deviation.

**Fig. 4.**
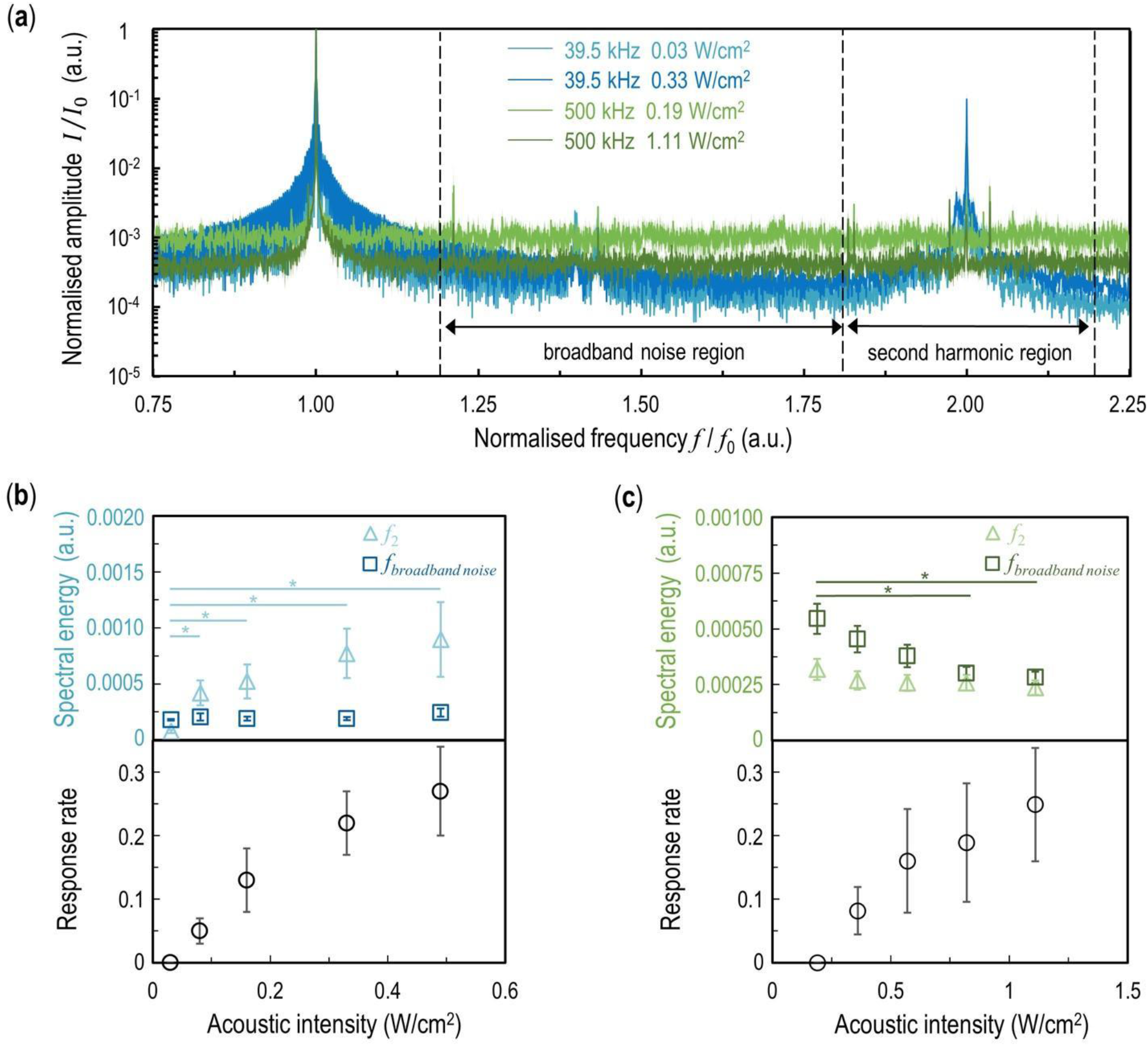
Cavitation occurrence during ultrasound stimulation. **(a)** Representative acoustic spectra of 39.5 kHz ultrasound stimulation with 0.03 and 0.33 W/cm^2^ acoustic intensity (blue) and 500 kHz ultrasound stimulation with 0.19 and 1.11 W/cm^2^ acoustic intensity (green). **(b)** Broadband noise, the second harmonic energy and response rate as a function of acoustic intensity for 39.5 kHz and **(c)** 500 kHz acoustic frequency (N = 6 dishes). Error bars and * represent standard deviation and p < 0.05, respectively.

### 3.2 39.5 kHz and 500 kHz ultrasound activate cortical cell by a different mechanical effect

Numerous in vivo studies have indicated that cavitation is involved in ultrasound stimulation from the observation of enhanced responses at lower frequencies of the sub-MHz band [42]. We assessed cavitation-triggered cell responses in cortical cells by measuring the occurrence of cavitation. Second harmonic spectral energy and broadband noise energy were measured from acoustic signals to evaluate the degree of cavitation. The second harmonic energy increased when cortical cells began to show responses to 39.5 kHz ultrasound stimulation **(Fig. 4a)**. The increase in cell response was found to be correlated with the increase of the second harmonic energy at 39.5 kHz **(Fig. 4b)**. These results indicate that stable cavitation activity plays a vital role in evoking cell responses to 39.5 kHz ultrasound neurostimulation. Increased cell responses were also observed at 500 kHz neurostimulation as ultrasound intensity was increased, whereas the second harmonic energy was not associated with increased cell responses as seen in 39.5 kHz neurostimulation **(Fig. 4c)**. The relationship between the second harmonic energy and cell responses was found to differ between 39.5 kHz and 500 kHz. Moreover, the increase in amplitude at the fundamental frequency was greater than other spectral components at 500 kHz, resulting in a decreasing trend in normalised broadband noise energy **(Fig. 4c)**.

Physical mechanisms, but not cavitation, were responsible for evoked responses at 500 kHz, albeit 500 kHz is within the sub-MHz range. This result indicates that the mechanical force from ultrasound is involved in neurostimulation. In light of previously reported studies on ultrasound neurostimulation [28], [43], [44], we consider that neurons are stimulated by the radiation force from 500 kHz ultrasound exposure. The calculated radiation force required for cell responses to 500 kHz ultrasound was ∼10^−4^ N/m^3^, whereas cells started to show response below this value for 39.5 kHz ultrasound neurostimulation. Cortical cells were subjected to a large radiation force during exposure to 500 kHz ultrasound, which evoked responses without cavitation activity, consistent with previous studies using high acoustic frequency range of the sub-MHz band [45].

Our results demonstrate that the physical mechanism underlying ultrasound neurostimulation is dependent on frequency; cavitation stimulates cells at low-frequency ranges and radiation force stimulates cells in high-frequency ranges. These mechanical effects could potentially perturb cellular structures [46] including the lipid bilayer [43] to activate mechanosensitive ion channels [42], [47], [48]. Measurement of this displacement would further elucidate the physical process of ultrasound neurostimulation. Furthermore, experiments performed between and beyond the frequency range from 39.5 kHz to 500 kHz can contribute to the search for finer demarcation of dominated mechanism transition.

## 4. Conclusions

We stimulated *in vitro* primary cortical cells with 39.5 kHz and 500 kHz acoustic frequencies and observed acoustic frequency-dependent responses, using fluorescence imaging of calcium responses, with acoustic signal recording. The response rate to 39.5 kHz ultrasound stimulation increased as acoustic signals at the second harmonic frequency component increased. In 500 kHz ultrasound stimulation, the second harmonic frequency did not show an increasing trend in response to cells’ increased activity. Accordingly, the physical effect of ultrasound neuromodulation is dependent on acoustic frequency where cavitation and radiation force are the dominant factors in low and high-frequency ranges, respectively. The results of the present study provide important insights into the acoustic frequency-dependent mechanism of ultrasound neuromodulation, where different physical effects evoke responses to low and high sub-MHz acoustic frequency ranges.

## Acknowledgements

This work was supported in part by JSPS KAKENHI Grant Number 19K20662 and 20K20642. A. I. was partly supported by JST PRESTO (JPMJPR1902).

## Competing interests

The authors declare no competing interests.

## Supplementary Data

**Supplementary Video 1**. Fluorescence change in cortical cells recorded at a frame rate of 60 fps during ultrasound stimulation (39.5 kHz, 0.33 W/cm^2^, 41.7 ms pulse duration) after image differencing.

**Supplementary Video 2**. Fluorescence change in cortical cells recorded at a frame rate of 60 fps during ultrasound stimulation (500 kHz, 1.11 W/cm^2^, 41.7 ms pulse duration) after image differencing.

## Notes

### Competing Interest Statement

The authors have declared no competing interest.

